# Genome announcement and analysis of five new mycobacteriophages belonging to the F, G, K, and P clusters

**DOI:** 10.1101/2024.02.15.580488

**Authors:** Ritu Arora, Ritam Das, Kanika Nadar, Urmi Bajpai

## Abstract

Mycobacterial infections are accountable for some of the most challenging to treat diseases in humans and animals. With the rise in drug-resistant clinical isolates of *Mycobacterium spp*., phage therapy is emerging as a promising alternative to antibiotic therapy. Isolation of genetically diverse phages that can infect pathogenic mycobacterium species is an important exercise. Here, we present the complete genome sequences of five mycobacteriophages isolated from environmental samples using *Mycobacterium smegmatis* Mc^2^ 155 as the bacterial host. These phages belong to different clusters, including F, G, K and P and contribute to the diversity of sequenced mycobacteriophages.

## ANNOUNCEMENT

*Mycobacterium smegmatis* is used as a surrogate for *Mycobacterium tuberculosis* and other non-tubercular mycobacteria, such as *M. fortuitum* and *M. abscessus* and phages infecting it have the potential to be useful therapeutically (*Shiloh et al*., 2010; *Hatfull et al*., 2022). Therefore, *M. smegmatis* serves as an efficient model organism for phage isolation. Genomes of about 18% (2383) of the total isolated mycobacteriophages (13290) (https://phagesdb.org/) have been sequenced and annotated (https://phagesdb.org/) and provide insights into the diversity, structure, and evolution of mycobacteriophages. Phage genome characterization advances our understanding of their biology and clinical importance, which can be applied in phage therapy and other applications.

To add to the existing repertoire, genome analysis of five new mycobacteriophages belonging to different clusters/sub-clusters is reported here. Environmental samples were collected from various regions of Delhi-NCR and Mumbai, India. The phages were isolated, purified and amplified on *M. smegmatis* Mc^2^ 155 as the host, using the double agar overlay method previously described by *Das et al., 2023*. Each phage was subjected to three rounds of purification and amplified to a titre >10^10^ pfu/ml. The high-titre lysates were used for genomic DNA isolation using the PCI method described previously by *Sinha et al., 2020*. Whole genome sequencing was outsourced, and the genomes were sequenced using the Illumina platform (150×2 PE chemistry). The WGS data was subjected to quality check using FastQC. The final assembled genomes were obtained as FASTA files, which were annotated using DNA master (https://phagesdb.org/DNAMaster/) and RAST-tk *(Brettin et al., 2015)* for predicting the CDS region. Blastp *(Altschul et al., 1990)*, HMMER *(Finn et al., 2011)* and InterProScan *(Jones et al., 2014)* tools were used for functional annotation. t-RNA was determined using ARAGORN *(Laslett et al., 2004)* and tRNAscan-SE *(Lowe et al., 2016)*. All programs were executed using default parameters. The sequenced and annotated genomes were submitted to GenBank (https://www.ncbi.nlm.nih.gov/genbank/). Table 1 describes all the five genomes and their features.

**Table 1:** Genomic characterization and features of five mycobacteriophages.

| **Phage**<br>**Name** | **Genbank**<br>**Accession**<br>**number** | **Cluster** | **Genome**<br>**size** | **G+C**<br>**content**<br>**(%)** | **ORFs** | **No. of**<br>**tRNAs** |
| --- | --- | --- | --- | --- | --- | --- |
| **KanikaMF1** | OQ985197.1 | P1 | 47372 bp | 67.2 | 78 | 0 |
| **Nanz197** | OQ555808.1 | K1 | 62663 bp | 66.5 | 98 | 1 |
| **RitSun** | MZ343158.1 | F2 | 55975 bp | 60.8 | 111 | 1 |
| **RitVan** | OQ201593.1 | G1 | 42068 bp | 66.8 | 62 | 0 |
| **Ritam007** | OP490271.1 | F4 | 55105 bp | 61.4 | 89 | 0 |

Phylogenetic analysis and sequence alignment were performed with the whole genome sequences of these five phages using the ViPTree software *(Nishimura et al., 2017)*. The analysis was performed against a curated database of several viral whole genome sequences belonging to different families and host groups. The five mycobacteriophages in this study were found to share phylogeny with the viruses infecting Actinomycetota, as expected. *RitVan* shared the same clade with Grizzly (MH779505), another mycobacteriophage from the G1 sub-cluster. *Nanz197* was found to have a separate clade, which appears to have diverged from the mycobacteriophage Murucutumbu (KM677211) and LaterM (MG962371) belonging to the K1 sub-cluster. *KanikaMF1* shares the clade with Malithi (KP027200), a P1 mycobacteriophage (Figure 1A).

**Figure 1:**
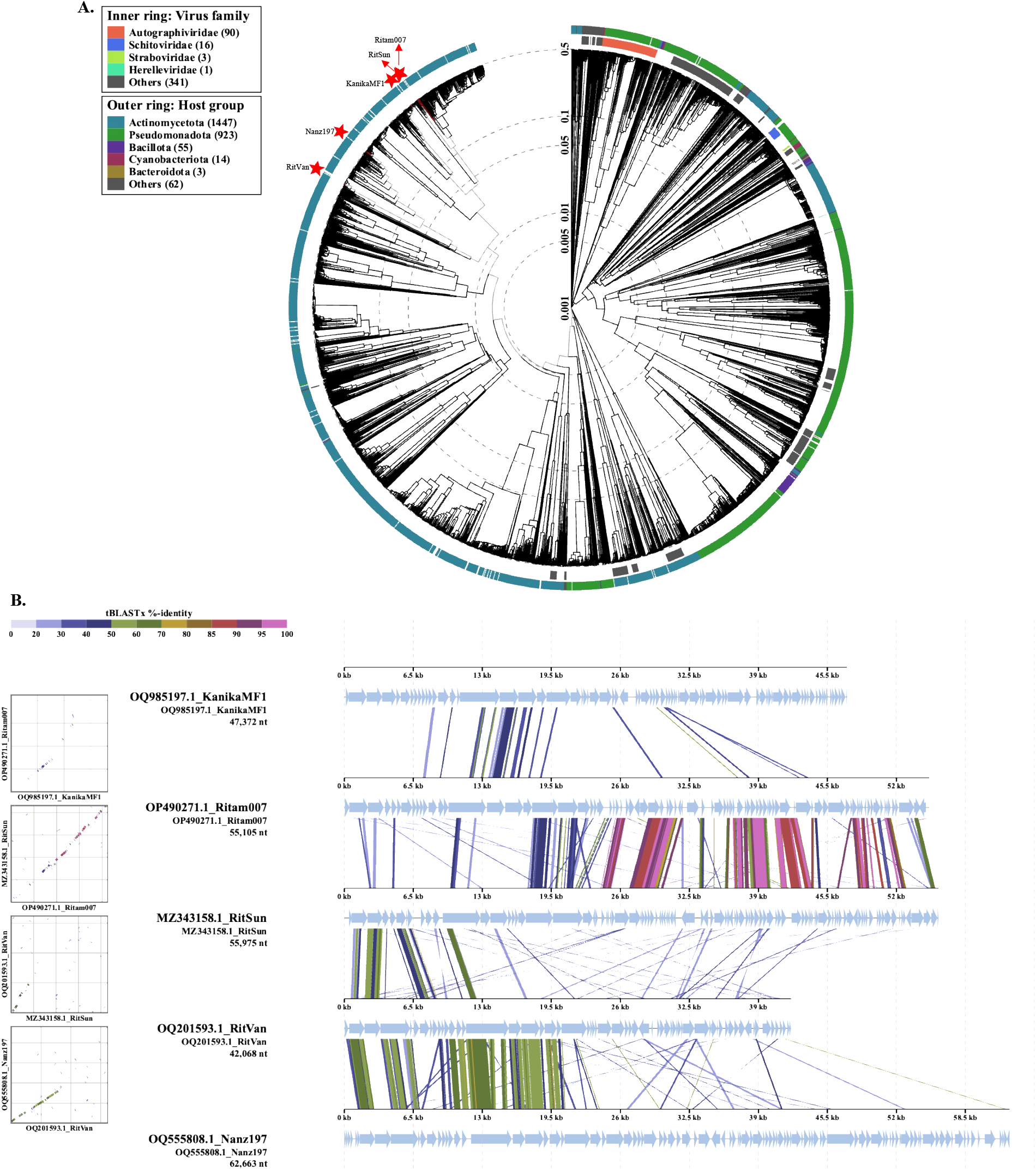
Sequence analysis of five mycobacteriophages discovered in this study. **A)** Phylogenetic analysis of the genomes demonstrated their shared ancestry with other phages infecting the host group Actinomycetota. The reference genome list had 2580 sequences from various virus families and was processed using VipTree software. B) Dot plot and sequence alignment of the five genomes indicate sequence similarities and differences. *Ritam007* and *RitSun* from the same cluster showrelatively high sequence identity, while *Nanz197* and *RitVan* share high sequence identity only in the anterior region of the genome.

According to our dot plot and sequence alignment analysis, *Ritam007* and *RitSun* are the most similar phages of the five studied (Figure 1B). They both are F-cluster mycobacteriophages that diverge in the phylogenetic analysis, forming their separate clades with their respective sub-cluster homologs (Figure 1A). *RitSun* was found to share a similar clade with mycobacterium phage Che9d (AY129336) from the F2 sub-cluster. *Ritam007*, like *Nanz197*, also formed its separate clade and appears to have diverged from the F4 sub-cluster mycobacteriophages Renaud18 (MH651187), ThetaBob (MK977709), and TChen (MH077585). Sequence alignment and dot plot analysis also demonstrated unique features in these genomes. As discussed above, *Ritam007* and *RitSun* belong to the same cluster (F) and show a high percentage of sequence identity.

Interestingly, *RitVan* (G1) and *Nanz197* (K1), even though distantly related, share a unique stretch of approximately 20 kb nucleotide sequence in the anterior region, demonstrating high sequence identity (Figure 1B). This region in both the genomes encodes for the structural proteins of the phage.

## DATA AVAILABILITY

The genome sequences of KanikaMF1, Nanz197, RitSun, RitVan and Ritam007 are available at GenBank under the accession number OQ985197.1, OQ555808.1, MZ343158.1, OQ201593.1, OP490271.1 respectively.

## ACKNOWLEDGEMENTS

We thank the Indian Council of Medical Research (ICMR), New Delhi, India, for supporting the project (5/8/5/38/2019-ECD-I). RA received the Senior Research Fellowship (SRF) from the University Grants Commission (UGC), New Delhi, India; RD received the ELITE fellowship from Acharya Narendra Dev College (ANDC), University of Delhi; KN received the Innovation in Science Pursuit for Inspired Research (INSPIRE) fellowship from the Department of Science and Technology (DST), India. We thank Acharya Narendra Dev College (ANDC), University of Delhi, for providing the infrastructure.

## REFERENCES

Altschul, S. F., Gish, W., Miller, W., Myers, E. W., & Lipman, D. J. (1990). Basic local alignment search tool. Journal of molecular biology, 215(3), 403–410.

Brettin, T., Davis, J. J., Disz, T., Edwards, R. A., Gerdes, S., Olsen, G. J., …& Xia, F. (2015). RASTtk: a modular and extensible implementation of the RAST algorithm for building custom annotation pipelines and annotating batches of genomes. Scientific reports, 5(1), 1–6.

Das, R., Arora, R., Nadar, K., Saroj, S., Singh, A. K., Patil, S. A., … & Bajpai, U. (2023). Insights into the genomic features, lifestyle and therapeutic potential of B1 sub-cluster mycobacteriophages. bioRxiv, 2023–05.

Finn, R. D., Clements, J., & Eddy, S. R. (2011). HMMER web server: interactive sequence similarity searching. Nucleic acids research, 39(suppl_2), W29–W37.

Hatfull, G. F. (2022). Mycobacteriophages: from petri dish to patient. PLoS Pathogens, 18(7), e1010602. https://phagesdb.org/ https://phagesdb.org/DNAMaster/ https://www.ncbi.nlm.nih.gov/genbank/

Jones, P., Binns, D., Chang, H. Y., Fraser, M., Li, W., McAnulla, C., … & Hunter, S. (2014). InterProScan 5: genome-scale protein function classification. Bioinformatics, 30(9), 1236–1240.

Laslett, D., & Canback, B. (2004). ARAGORN, a program to detect tRNA genes and tmRNA genes in nucleotide sequences. Nucleic acids research, 32(1), 11–16.

Lowe, T. M., & Chan, P. P. (2016). tRNAscan-SE On-line: integrating search and context for analysis of transfer RNA genes. Nucleic acids research, 44(W1), W54–W57.

Nishimura, Y., Yoshida, T., Kuronishi, M., Uehara, H., Ogata, H., & Goto, S. (2017). ViPTree: the viral proteomic tree server. Bioinformatics, 33(15), 2379–2380.

Shiloh, M. U., & Champion, P. A. D. (2010). To catch a killer. What can mycobacterial models teach us about Mycobacterium tuberculosis pathogenesis?. Current opinion in microbiology, 13(1), 86–92.

Sinha, A., Eniyan, K., Manohar, P., Ramesh, N., & Bajpai, U. (2020). Characterization and genome analysis of B1 sub-cluster mycobacteriophage PDRPxv. Virus research, 279, 197884.

